# α7 nicotinic acetylcholine receptors are necessary for basal forebrain activation to increase expression of the nerve growth factor receptor TrkA

**DOI:** 10.1101/2024.03.01.582932

**Authors:** Jacob Kumro, Ashutosh Tripathi, Alvin V. Terry, Anilkumar Pillai, David T Blake

## Abstract

Activation of the basal forebrain leads to increases in the expression of the nerve growth factor receptor, Tropomyosin receptor kinase A (TrkA) and decreases in expression of the beta amyloid cleavage enzyme 1 (BACE1) in the cerebral cortex of both sexes of 5xFAD mice. The studies described in this report were designed to determine if these changes were dependent on acetylcholine receptors. Mice were stimulated unilaterally in the basal forebrain for two weeks. Animals were administered a cholinergic antagonist, or saline, 30 minutes prior to stimulation. Animals administered saline exhibited significant increases in TrkA expression and decreases in BACE1 in the stimulated hemisphere relative to the unstimulated. While both nonselective nicotinic and muscarinic acetylcholine receptor blockade attenuated the BACE1 decline, only the nicotinic receptor antagonism blocked the TrkA increase. Next, we applied selective nicotinic antagonists, and the α7 antagonist blocked the TrkA increases, but the α4β2 antagonist did not. BACE1 declines were not blocked by either intervention. Mice with a loxP conditional knockout of the gene for the α7 nicotinic receptor were also employed in these studies. Animals were either stimulated bilaterally for two weeks, or left unstimulated. With or without stimulation, the expression of TrkA receptors was lower in the cortical region with the α7 nicotinic receptor knockdown. We thus conclude that α7 nicotinic receptor activation is necessary for normal expression of TrkA and increases caused by basal forebrain activation, while BACE1 declines caused by stimulation have dependency on a broader array of receptor subtypes.

## INTRODUCTION

Despite decades of research, the current clinical therapies for Alzheimer’s disease (AD) still primarily focus on increasing synaptic concentrations of acetylcholine in the brain to improve cognitive deficits. This treatment is based on the cholinergic hypothesis of AD which links the volume loss and cytological changes in the subpallidal basal forebrain, the source of neocortical cholinergic afferents (Mesulam & Van Hoesen, 1976; Schmitz & Nathan Spreng, 2016; Whitehouse et al., 1982) as well as a reduction in presynaptic cholinergic markers in the cortex (Bowen et al., 1976; Davies & Maloney, 1976) to subsequent memory impairment and beta-amyloid accumulation in AD brains. Further research demonstrated the importance of nerve growth factor (NGF) signaling during synaptic activity to maintain cholinergic synaptic integrity specifically in the cortex and hippocampus (Cuello, 2013; Cuello et al., 2007, 2010) which is also relevant to the pathophysiology of AD. While mature NGF, mNGF, promotes cholinergic neuron survival through interactions with its TrkA receptor, the precursor of NGF, proNGF, has higher affinity for p75NTR and is associated with apoptosis (Lee et al., 2001). AD brains demonstrate higher concentrations of proNGF and lower concentrations of TrkA which supports NGF signaling imbalance as a potential contributor to AD pathology and dementia (Counts et al., 2004; Iulita & Cuello, 2014; Mufson et al., 2000). Clinical applications of exogenous (intraventricular) administration of NGF to the cerebrum were associated with off-target side effects associated with pain signaling (Eriksdotter Jönhagen et al., 1998), indicating that approaches designed to more selectively enhance NGF signaling in the appropriate brain region (i.e., basal forebrain cholinergic pathway) are needed.

Electrical deep brain stimulation (DBS) of the basal forebrain has been proposed as a means of maintaining trophic support and increasing NGF release (Hotta et al., 2007, 2009). We have shown that basal forebrain DBS increases TrkA receptor levels in the frontal cortex and decreases beta-amyloid cleaving enzyme 1 (BACE1) in an Alzheimer’s mouse model (Kumro et al., 2023). We observed similar effects of DBS on TrkB, the high affinity receptor for brain-derived neurotrophic growth factor (BDNF). Importantly BDNF and its receptor kinase TrkB, which is also reduced in AD, is transported as a complex (similar to NGF/TrkA) in a retrograde manner, from the axon terminal to the nucleus in the soma, to provide trophic support to many neuronal populations (Iulita et al., 2017). DBS in other brain regions also increases BDNF in various neural circuits (Dandekar et al., 2018; Do-Monte et al., 2013; Sun et al., 2022).

One interesting question that has yet to be fully addressed is whether the positive effects of DBS on neurotrophin receptors noted above are dependent on cholinergic receptors and if so, if the effects are receptor subtype selective.

Several early observations would appear to suggest that nicotinic acetylcholine receptors might be especially important. The initial reports of neuroprotective effects of the tobacco alkaloid nicotine both *in vitro* and *in vivo* models of neural toxicity (Belluardo et al., 2000; O’Neill et al., 2002) as well protracted positive effects on memory function in monkeys for up to 24 h after a single μg/kg dose of nicotine (Buccafusco & Jackson, 1991; Terry et al., 1993) led us to hypothesize that neurotrophins or their receptors might be involved. This hypothesis was initially tested *in vitro* in differentiated PC-12 cells and the results indicated that exposure to nicotine enhanced the expression of NGF-receptors (Terry & Clarke, 1994). Subsequent *in vitro* and *ex vivo* experiments with selective antibodies indicated the effects of nicotine on NGF receptors were selective for high affinity TrkA receptors as opposed to low affinity p75NTR receptors, and further, using selective nicotinic receptor antagonists (e.g., methyllycaconitine), that the effects were dependent on α7 nicotinic receptors (Hernandez & Terry, 2005; Jonnala et al., 2002).

It is also important to note that anti-NGF and anti-BDNF antibodies each reduce α7 nicotinic receptor density in the hippocampus (Kawai et al., 2002). Moreover, the basal forebrain, including the subpallidal region, is one of the most densely populated areas of the brain with α7 receptors which have high affinity for beta-amyloid resulting in its antagonism (Hernandez et al., 2010; H.-Y. Wang et al., 2002).

In the experiments described in this report, we, therefore, used unilateral, intermittent basal forebrain DBS while applying selective and non-selective cholinergic receptor antagonists to determine the dependence of the neurotrophin receptor expression changes on specific acetylcholine receptor subtypes (Kumro et al., 2023). We used the 5xFAD model to promote BACE1 baseline expression, as prior work found low levels in wildtype mice.

## Materials and Methods

### Test Subjects

B6SJL-Tg(APPSwFlLon,PSEN1*M146L*L286V)6799Vas/Mmjax or 5XFAD as well as B6(Cg)-*Chrna7*^*tm1.1Ehs*^/YakelJ or α7nAChR^flox^ mice of both sexes were used for this study. Transgenic 5XFAD mice were identified by PCR using a sense primer (5’-CTA CAG CCC CTC TCC AAG GTT TAT AG-3’) and antisense primers (5’-AAG CTA GCT GCA GTA ACG CCA TTT-3’) and (5’-ACC TGC ATG TGA ACC CAG TAT TCT ATC-3’). Heterozygote α7nAChR^flox^ mice were identified by PCR using a sense primer (5’-GTC CCT CTG CTG GTA TTT GC-3’) and antisense primers (5’-GAA CAA GTC AGA TAA GAA CCT-3’) and (5’-GCC CAA TTC CGA TCA TAT TC-3’). PCR products were separated on a 2% agarose gel, and a DNA ladder confirmed correct band size. Mice were housed for four months before all experiments. Mice were housed in groups of 5 per cage, in a temperature and humidity-controlled room on a 12-hour light and dark cycle (06:00-18:00). Animals were given free access to food (Teklad Rodent Diet 8604 pellets, Harlan, Madison, WI) and water. The Institutional Animal Care and Use Committee of Augusta University approved of all protocols and procedures used in this study. Under the guidance of the 8th Edition of the National Institute of Health Guide for Care and Use of Laboratory Animals (2011), measures were taken to minimize pain and discomfort throughout the study.

### Electrode Fabrication

Stimulation electrodes were custom-made based on previously published specifications(McCairn & Turner, 2009). Stainless steel 30 ga. hypodermic tubing (MicroGroup ®, Medway, MA) was cut to a length of 2.8mm using a Dremel ® rotary saw. A 26G (Precision Glide ®) needle was twisted into each end of the tubing to re-establish their patency and a tungsten wire reamed out any other debris. Teflon coated platinum-iridium wire with an outer diameter of 139.7μm (A-M Systems, Seattle, WA) was cut to 20mm in length and threaded through the hypodermic tubing. Iris scissors grasped and unsheathed 0.5 mm of the Pt-ir wire to create the conductive tip. Polyimide tubing was cut to a 3.8mm length and slid onto the hypodermic tubing and wire so that the 1.0mm excess polyimide tubing was left at the tail end of the electrode. The stimulation tip was adjusted in the hypodermic tubing exposing the 0.5mm of uncoated wire. Using the included brush, cyanoacrylate (Krazy ®) was applied to the excess polyimide opening until glue was drawn into the area creating a 4.8mm long electrode from the stimulation tip to end of polyimide tubing. A butane torch uncoated about 3mm of the tail-end wire.

### Surgical Procedures

#### Stimulation Electrode and Headcap Implantation

Surgery was performed on a sterile field under 1.5% isoflurane anesthesia. The mouse was placed in a dual arm, rodent stereotaxic instrument (Stoelting ®, Wood Dale, IL) fitted with an anesthesia mask and heating pad set to 37°C. Stereotaxic measurements were made in reference to the bregma landmark (−0.6mm AP or caudal, 1.7mm ML or lateral) to target sites for electrode implantation. A 0.5 mm hole was drilled at target sites using a carbide bur (HM1-005-FG, Meisinger, Centennial, CO) and dental drill (Volvere GX, Brasseler, Savannah, GA) until the cranium was breached. An electrode was loaded into a guide tube positioned above the target site and pushed into the cerebrum with a stylus until the back of the electrode became flush with the cortical surface to ensure achievement of 4.8mm depth. Cyanoacrylate was placed at the target site to stabilize the electrode and seal off exposed tissue. The procedure was repeated for implantation of the second electrode in the contralateral hemisphere. All animals in the pharmacology studies were stimulated equally unilaterally. With the same burr, a hole was made at each temporal and frontal bone and stainless steel screws (Antrin, Fallbrook, CA) were hand screwed two rotations. Before implanting the final bone screw in the left frontal bone, a 20mm length of Pt/Ir wire with 3mm of insulation removed at each end was used. One end was bent with forceps and then slid into the bone screw hole so that the tip resided between the top of the cortex and bottom of the skull. A layer of cyanoacrylate was applied to all screws and electrode sites and left until dry. Dental cement was applied to the top of the skull with the periosteal elevators. A type B micro-USB receptacle (part no. 84×0185, Newark®, Chicago, IL), with the 2^nd^ and 4^th^ connector pins removed, was fixed to one of the stereotaxic arms and positioned over the skull. The exposed wire from the bone screw was soldered to the middle grounding pin of the micro-USB port while the electrode wires were soldered to the remaining two outer pins. A thin coat of dental cement was applied to the soldered pins to ensure firm connection. The micro-USB receptacle was dismounted from the stereotactic arm and positioned just above the top of the bone screws. A larger stainless steel screw (MX-M025, Small Parts Inc., Logansport, IN) was adhered to the side of the micro-USB receptacle that would serve as a handle when connecting mice to the stimulating computer. Dental cement was applied to fill in all areas between the base of the skull and base of the micro-USB receptacle to create the stimulating headcap. A micro-USB receptacle was placed into the top of the headcap to prevent cage-mates from chewing on the internal connector. The mouse was returned to the cage and monitored until conscious and consuming water and food.

Lesions were created in pilot animals until targeting was consistent. Lesions were created with a 30 μA DC current, electrode negative, for 10 seconds which generated a clearly visible mark on perfused and unstained sections (Kumro et al., 2023).

#### Viral Transduction

Surgery was performed on a sterile table under 1.5% isoflurane anesthesia. Artificial tears ophthalmic ointment was applied to both eyes. A shaved mouse was transferred to the rodent stereotaxic instrument. A 3mm incision was made with surgical scissors at an approximated location to expose the skull near 2mm lateral and 2mm posterior to bregma. A small hole was carefully made (−2.0mm AP, 2.0 mm ML from bregma) over the somatosensory cortex. A micropipette was slowly inserted to a depth of 700μm. α7nAChR^flox^ mice were injected with 500nL of AAV9-Cre-GFP (Vector Biolabs) at 100nL/min using a micropump (WPI). After a 5-min wait period, the micropipette was slowly withdrawn. The scalp incision was sutured closed. The procedure was repeated for the contralateral side and injection of 500nL of sterile 0.9% saline was made in a similar fashion. Mice were returned to the home cage, and the viral vector incubated for 2-wks. To confirm vector expression, mice to undergo stimulation were anesthetized and prepped as stated above. The scalp was re-incised to expose the injection sight, and the presence of fluorescence was checked using confocal epifluorescence microscopy fitted with a FITC filter (emission of 513-556nm, excitation of 467-498nm). Mice demonstrating positive fluorescence were then implanted with microelectrodes and a headcap as detailed above.

### Brain Stimulation

#### Electrode Impedance Measurement

The individual impedances of each electrode had to be calculated so that the stimulation voltages could be adjusted to deliver 100 μA of current to each electrode tip. Because previous pilot studies demonstrated a consistent daily impedance, impedances were measured at the beginning of each week. Impedance was measured using custom programmed software that triggered a stimulus isolator (A385 World Precision Instruments, Sarasota, FL) to produce a 100 μA current square wave and then use an oscilloscope (TDS 2012C Tektronix, Beaverton, OR) to measure the voltage change. The same method was applied to the left and right electrodes of each mouse head cap. Impedances averaged 8-14 kOhm. Resistors were added to the stimulation cord circuit path as necessary in order to stimulate multiple mouse electrodes at the same time with a uniform voltage output. Typically, stimulation was delivered at close to 1 volt peak per phase across an impedance of 10 kΩ

#### Mouse Stimulation

Mice were brought to the stimulation room and allowed 30-min of acclimation before beginning stimulation. Remaining in their home cages, the large screw adhered to the side of the micro-USB receptacle was secured with hemostats while a micro-USB cable was plugged into each mouse headcap. The mouse cage lids were returned ajar so that mice had continued access to food and water. All micro-USB charging cables were threaded through a pulley system equipped with a counter balance and suspended from the ceiling to remove any upward or downward forces on the headcaps during stimulation and allowing for better mouse mobility. Tantalum capacitors (10μF 25v, Kemet, Fort Lauderdale, FL) were soldered to each stimulating cable end to block net charge transfer. These capacitors and the cable’s grounding wires were connected to a multiple functional I/O device (USB-6211, National Instruments, Austin, TX) that served as a voltage generator creating the voltage-dependent stimulation pulses signaled by custom programmed software. Stimulation was delivered with biphasic, negative first (cathodal), unipolar 100 μA pulses with 100 μS per phase. 60 pulses were delivered per second for 20-sec followed by 40-sec of no stimulation. Mice received 1-hr of unilateral stimulation 5-days each week in pharmacological studies, and 1 hr bilateral stimulation in α7nAChR^flox^ mouse studies. We estimate the activating function radius for 100 μA stimulation as 315 μm using prior work (McIntyre et al., 2004; Murasugi et al., 1993; Stoney et al., 1968). The stimulation typically evoked a brief voltage change of 1 volt(Rozman et al., 2000), safely within limits for platinum for brief pulses(Rose & Robblee, 1990), and had charge densities well below limits for tissue damage(Shannon, 1992).

### Tissue Processing

#### Brain Extraction

One day after their last stimulation, mice were placed in a bell jar with isoflurane until loss of righting reflex and unresponsive to toe pinch. Grasping the base of the skull and tail, manual cervical dislocation was performed to euthanize the mouse. Surgical scissors transected through cervical tissue and spinal cord in order to isolate the head. A periosteal elevator removed the brain from the cranium and transferred it face down onto a sterile petri dish on a bed of crushed ice. A #11 scalpel made a coronal incision at the origin of the cranial nerve II white matter tracts. The cortical portion of the anterior section was removed with forceps and a scalpel. The cortical portion of the posterior brain section was reflected from the midline using a blunt probe to expose both hemispheres of the hippocampus which were resected with forceps. All frontal cortex and hippocampus tissue was separated by hemisphere and transferred to labeled 1.7 mL microcentrifuge tubes on ice and then stored at -80°C.

#### Tissue Homogenization

100 μl of 4°C radioimmunoprecipitation buffer (Sigma-Aldrich, St. Louis, MO) with 10 μl/ml protease inhibitor cocktail (AEBSF at 104 mM, Aprotinin at 80 μM, Bestatin at 4 mM, E-64 at 1.4 mM, Leupeptin at 2 mM and Pepstatin A at 1.5 mM, Sigma-Aldrich, P8340) and was added to each frozen brain section on ice. Samples received a 5-sec pulse from an ultrasonic homogenizer (Qsonica ®, Newtown, CT) while sitting in crushed ice. Homogenized samples underwent centrifugation at 15,000 g for 15-min at 4°C. Supernatants were transferred to pre-chilled microcentrifuge tubes and stored at -80°C until used for protein analysis.

### Protein Analysis

#### Western Blot Analysis

Protein concentrations of each sample were determined using the bicinchoninic acid (BCA) protein assay kit (Thermo Fisher Scientific, Waltham, MA) which used bovine serum albumin standards to create a best fit linear regression line based on protein concentrations using the 562nm absorbance mode of a microplate reader and its Gen5 Data Analysis program (Biotek, Winooski, VT). Equal amounts of protein were resolved in SDS-polyacrylamide gels and electrophoretically transferred to nitrocellulose membranes (Bio-Rad, Hercules, CA) at 4°C. Membranes were blocked for 1-hr in Tris-buffered saline containing Tween 20 (TBST; 10mM Tris-HCl, pH 8.0, 138 mM NaCl, 2.7 mM KCl, and 0.05% Tween 20) and 5% non-fat milk.

Primary antibodies used were anti-TrkA (1:1000, GTX132966, GeneTex ®, Irvine, CA), anti-BACE1 (1:1000, GTX103757, GeneTex ®, Irvine, CA) and anti-GAPDH (1:1000, D16H11, Cell signaling technology, Danvers, MA).

Primary antibodies were added to 10 ml of 5% non-fat milk and TBST solution and applied to the blocked membrane to incubate overnight on an orbital shaker at 4°C. The following day, membranes were washed three times with TBST for 5-min and then incubated for 1-hr with horseradish peroxidase-conjugated goat anti-rabbit IgG (ab6721, Abcam, Cambridge, MA). Membranes were washed three more times with TBST for 5-min.

Blots were developed using Pierce™ ECL Western Blotting Substrate (ThermoFisher, USA, Cat# 32106) at a sufficient volume to ensure that the blot was completely wet with the substrate and the blot didn’t become dry (0.1 ml/cm^2^). Membrane incubated with the substrate working solution for 1 minute. Blot was removed from ECL solution and placed in a plastic sheet protector or clear plastic wrap. An absorbent tissue removed excess liquid and carefully pressed out any bubbles from between the blot and the membrane protector. Then the blot was analyzed using The ChemiDoc XRS+ Gel Imaging System (BioRad, USA). The images were analyzed using Image Lab image acquisition and analysis software (BioRad, USA).

To avoid saturation and ensure linearity, 30μg of each sample were loaded in the gel. Study has shown that GAPDH showed linearity up to 30 μg of protein loaded. Along with this, gels were stained with Coomassie stain to ensure equal amount of protein loaded (Eaton et al., 2013; Fosang & Colbran, 2015).

For the different molecular weights, membranes were cut into strips (horizontally using protein ladder/marker), then probed individually with different antibodies. However, for very close molecular weights and/or for the GAPDH, membrane blots were stripped using western blot stripping buffer (Thermo Fisher, USA, cat# 21059). For stripping, blots were washed 5 min x 3 with TBST to remove chemiluminescent substrate, and then incubated in western blot stripping buffer for 5 to 15 minutes at RT. Then they were again washed 5 min x 3 with TBST, and then performed next immunoblot experiment. The original blots are provided in the supplementary file.

### Statistical Analysis

Statistical analyses were performed using GraphPad Prism 9 software. Two-tailed Student’s t-tests were used for group comparisons and differences between means of experimental groups were considered significant at the p < 0.05 level.

## Results

The first sets of experiments used systemic pharmacological antagonism administered 30 minutes prior to basal forebrain stimulation. The stimulation was delivered for one hour daily, and was unilateral. The stimulation delivered unipolar, biphasic, negative-first, pulses for 20 seconds of each minute, and 40 seconds off each minute. Electrode impedance was checked weekly, and voltage was altered to set currents at 100 μA per phase. After two weeks of such stimulation, tissue was harvested and frontal cortex was collected for protein expression analysis. The stimulation typically evoked a brief voltage change of 1 volt (Rozman et al., 2000), safely within limits for platinum for brief pulses (Rose & Robblee, 1990), and had charge densities well below limits for tissue damage (Shannon, 1992).

### Baseline changes in TrkA and BACE1 caused by stimulation

Five 5xFAD mice were used in the saline experiments. Experiments tested the null hypothesis that TrkA and BACE1 expression were unchanged by two weeks of one hour daily unilateral stimulation applied 30 minutes after saline IP administration. TrkA and BACE1 expression from Western blots were compared, as shown in Figure 1. A paired samples statistical approach was used to assess changes in TrkA and BACE1. Expression of TrkA rose with stimulation (t(5) = 3.27, p<0.03), and BACE1 expression declined (t(5) = -6.07, p<0.004), which rejects the null hypothesis.

**Figure 1.**
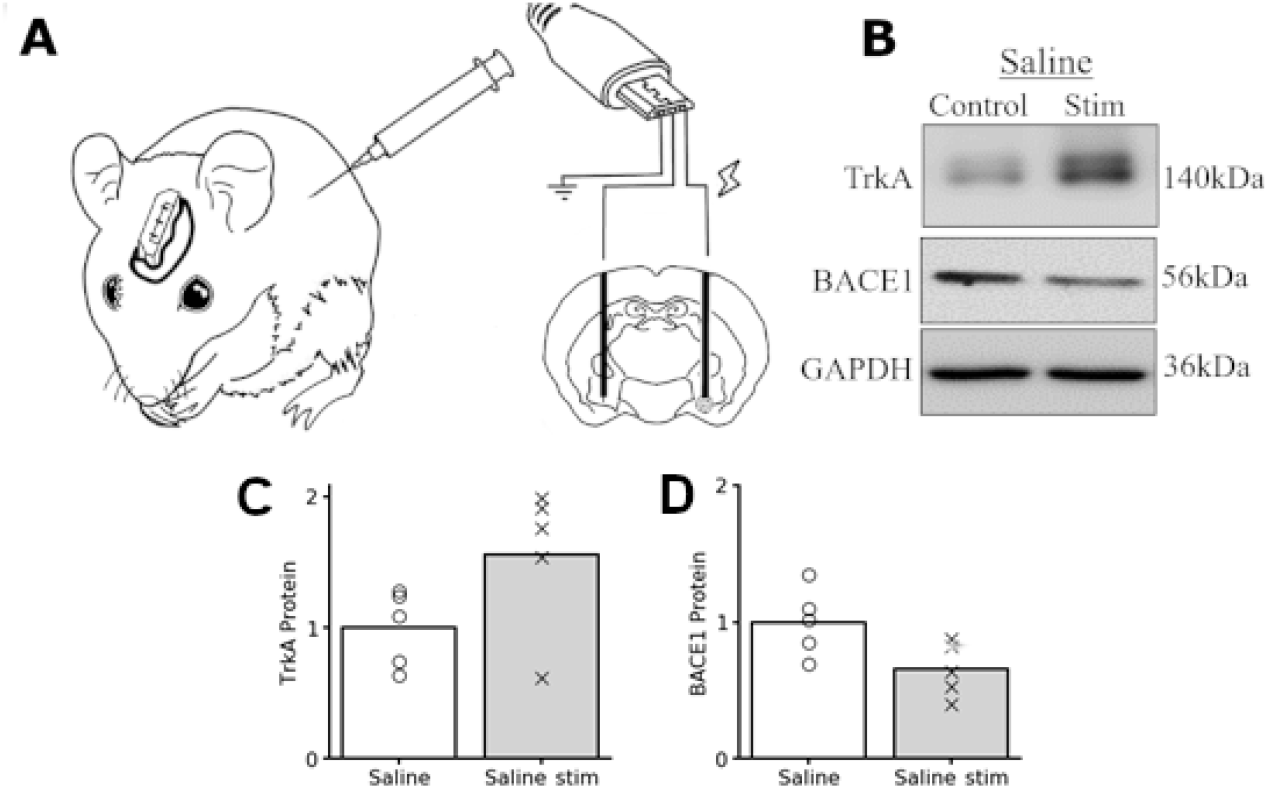
Protein expression changes in the control condition. A. Mouse with unilateral brain stimulation circuit B. Representative immunoblots C. Stimulated hemisphere is compared to the unstimulated hemisphere after saline administration for TrkA. D. Same, for BACE1. All data points for each bar are shown superimposed on the plot.

### Comparison of changes in TrkA and BACE1 caused by stimulation as impacted by muscarinic vs nicotinic antagonism

To narrow down potential acetylcholine receptors responsible for the TrkA and BACE1 changes in the saline experiments, non-selective antagonists were used in two groups of 5xFAD mice given either scopolamine or mecamylamine prior to stimulation to antagonize the muscarinic or nicotinic class of receptors respectively.

As shown in Figure 2, mice administered scopolamine demonstrated that TrkA trended higher in the stimulated hemisphere (t(4)=2.54, p<0.08), and BACE1 was only modestly lower in the stimulated hemisphere (t(4)=-1.33, p<0.27). However, mice administered mecamylamine prior to stimulation no longer demonstrated an increase in TrkA expression, but trended to decline modestly (t(5)=-0.9,p<0.4) and the decline in BACE1 was less pronounced compared to the non-stimulated hemisphere (t(5)=-1.41, p<0.23).

**Figure 2.**
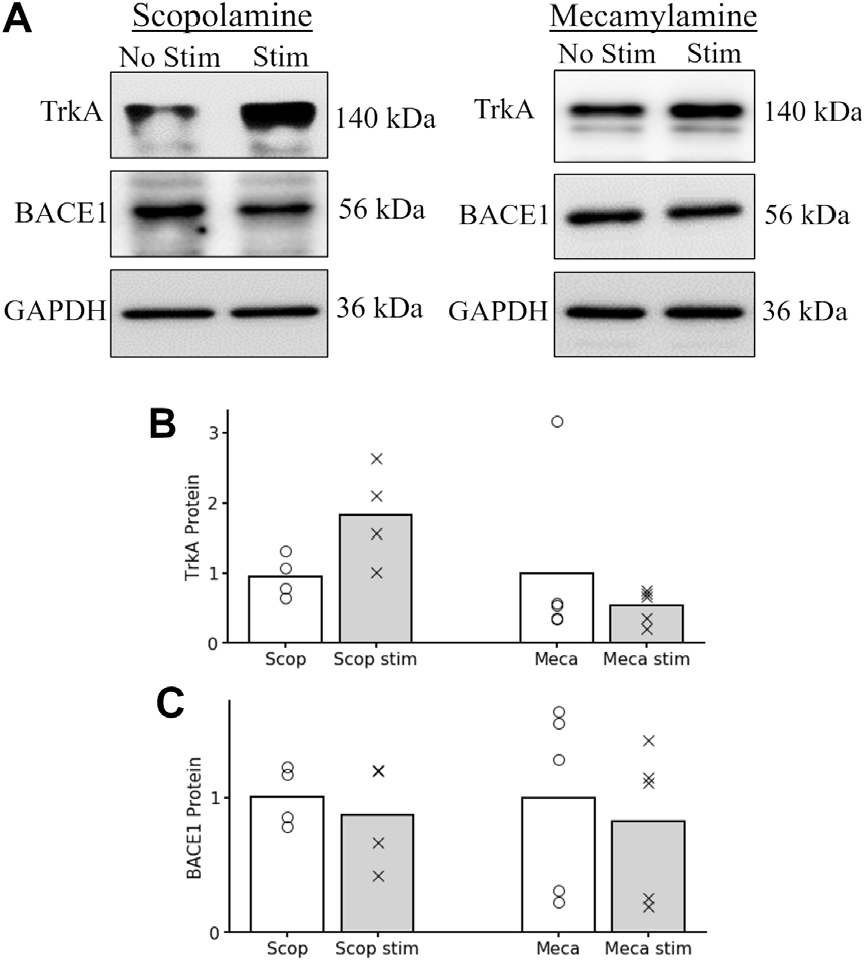
Protein expression changes compared under muscarinic or nicotinic antagonism. A. Representative immunoblots B. TrkA increases caused by stimulation as impacted by scopolamine or mecamylamine block. C. Same for BACE1 decreases.

### Comparison of changes in TrkA and BACE1 caused by stimulation as impacted by selective nicotinic antagonism

With non-selective nicotinic antagonism blunting stimulation effects, further selectivity was investigated with the two most common nicotinic receptors found in the cerebral cortex: α7 and α4β2. The α7-selective antagonist methyllycaconitine (MLA) was used in five mice of either sex, and the α4β2-selective antagonist dihydro-β-erythroidine (DHBE) was used in four animals of either sex.

The TrkA changes caused by stimulation trended up while using DHBE (t(4)=2.95, p<0.06) but did not trend up with MLA (t(5)=0.28, p<0.8). The BACE1 declines caused by stimulation trended down with either antagonist (DHBE t(4) = -2.97, p=0.06, MLA t(5) = -2.15, p=0.1).

We hypothesized that the cholinergic effects on TrkA were completely determined by α7 nicotinic receptor antagonism, but our data lacked adequate statistical power from low numbers. We regrouped the data to explore the impact of antagonizing each of the two nicotinic receptors. First, we tested the null hypothesis that basal forebrain activation without muscarinic or α4β2 receptor activation did not alter TrkA compared to the contralateral unstimulated hemisphere. Combining scopolamine and DHBE groups resulted in a stimulation dependent significant increase in TrkA, which rejects that null hypothesis (t(7)=4.08, p=0.005).

Next, we tested the hypothesis that basal forebrain activation with nonselective nicotinic and α7 specific antagonists did not alter TrkA expression across hemispheres. The group combining mecamylamine and MLA trended to have less TrkA expression on the stimulated hemisphere, which trends opposite to, and cannot reject, the null hypothesis (t(9)=-0.87, p<0.4).

Grouping the data with DHBE and mecamylamine in one group (both antagonize α4β2) did not result in a significant TrkA increase from stimulation t(8)=0.21, p<0.84, which was comparable to grouping scopolamine and MLA (do not antagonize α4β2) t(8)=1.91, p<0.09.

### The impact of neuron-specific knockdown of CHRNA7

The specificity of the TrkA stimulation increase for α7 nicotinic receptor antagonism motived the use of the α7nAChR^flox^ mouse. Seven mice of either sex were used in experiments without stimulation, and three were used in experiments with bilateral stimulation. A viral vector injection of 1 μL of AAV9-hSYN1-eGFP-2A-iCre-WPRE was performed in one hemisphere of the mice to express GFP and knockdown of CHRNA7, the α7 nicotinic receptor subunit. The hSYN1 promoter by design only alters expression in neurons. A matched volume of saline was infused contralaterally. After three weeks, expression was verified by qualitative fluorescence, and animals in the stimulation group were implanted with stimulating electrodes. Animals were stimulated bilaterally for one hour per day for two weeks, and then brains were harvested for protein analysis. In the group without stimulation, animals were left in the colony for two weeks prior to analysis. Only cortical tissue with GFP expression was sampled for the knockdown group, while a matched contralateral volume was sampled for the contralateral control.

As shown in Figure 4, without stimulation, the knockdown of the α7 receptor gene resulted in lowered TrkA than the contralateral tissue (t(7)=2.68, p<0.018). In the animals receiving stimulation, the ratio of TrkA in the control tissue to that in the knockdown tissue trended similarly (t(3)=1.16 p<0.18) but the number of samples used prevents a strong conclusion. Both groups show a more than two-fold reduction in TrkA in the hemisphere with a neural knockdown of CHRNA7, while our pharmacological experiments found the stimulation increased TrkA by nearly 50% in the saline control condition.

**Figure 3.**
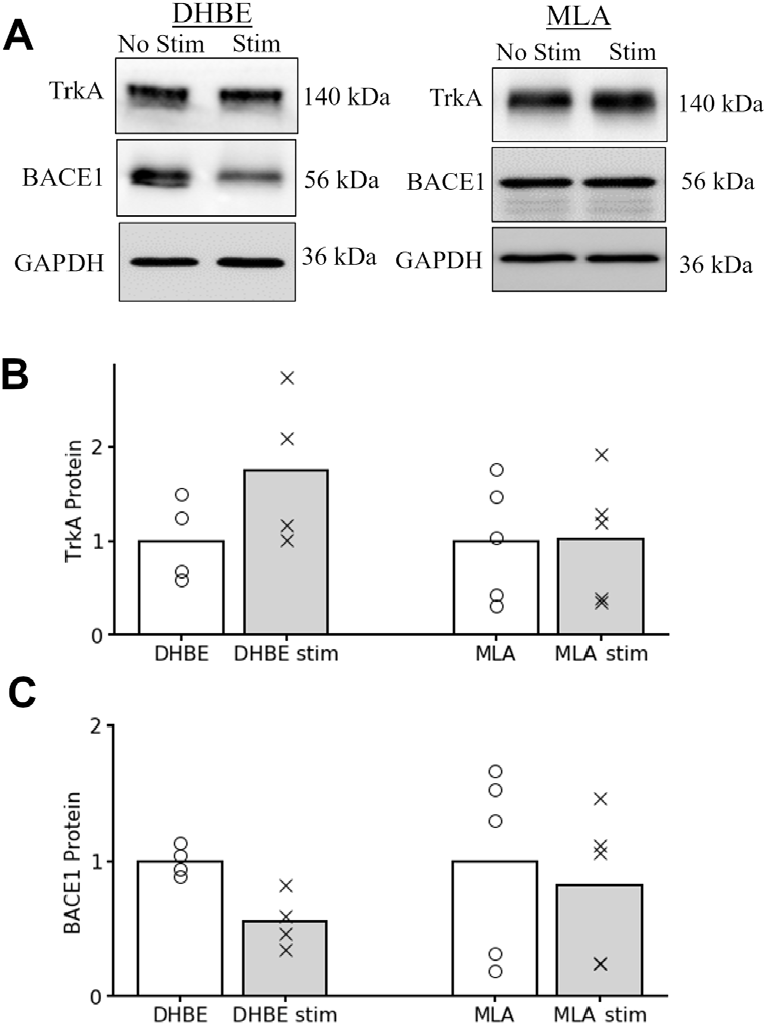
Protein expression changes compared under different selective nicotinic antagonism. A. Representative immunoblots B. TrkA increases caused by stimulation as impacted by administration of dihydro-β-erythroidine (DHBE) or methyllycaconitine (MLA) C. BACE1 decreases caused by stimulation impacted by DHBE or MLA.

**Figure 4.**
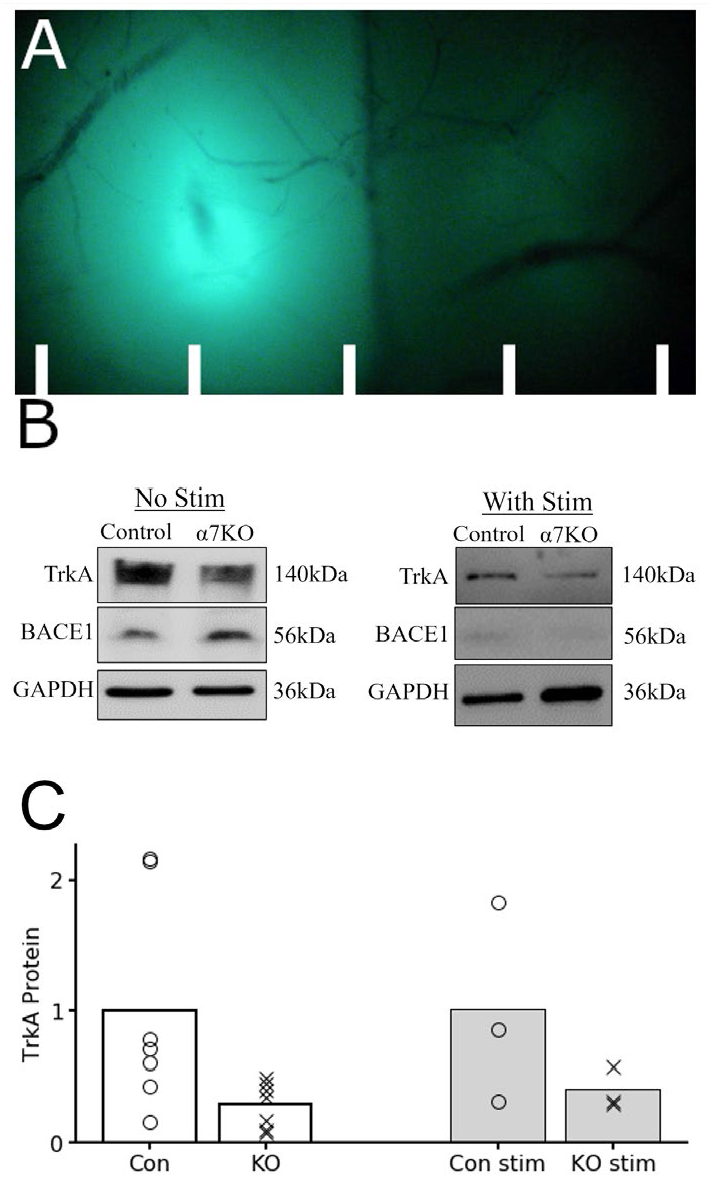
Effect of knockdown of Chrna7 on TrkA expression. A. Epifluorescence of both hemispheres shown, demarcated with 1mm lines. Both hemispheres were either stimulated or non-stimulated. The fluorescent tissue was sampled from the left, while a matched volume was sampled from the right. B. Representative immunoblots. C. The control vs knockout results are shown on the left, and the stimulated group on the right.

## Discussion

Previous findings demonstrated that unilateral, intermittent basal forebrain DBS increases expression of the nerve growth factor receptor TrkA(Kumro et al., 2023). Assays for other relevant TrkA related proteins such as NGF, proNGF, and phosphyrylated TrkA were inconsistent in comparison, perhaps because tissue harvest occurred 24 hours after the last stimulation, and the activated NGF pathway has a time course closer to 6 hours(Kaplan et al., 1991). Because acetylcholine release increases NGF and its receptor, we chose TrkA expression as the representative variable for the neurotrophic pathways that maintain cholinergic tone due to its longer half-life than NGF (Hotta et al., 2009; Jonnala et al., 2002). The NGF and TrkA interplay is paramount in the development of the cholinergic system. During this period, cholinergic neurons expressing TrkA produce growth cones and extend their axonal projections into tissues that produce NGF like the cortex (Gallo et al., 1997). After development, TrkA still remains important in the integrity and maintenance of this system as increased acetylcholine release increases axonal branching and cholinergic enzyme markers, and TrkA antagonism impairs these processes leading to axonal withdrawal (Debeir et al., 1999; Ginty & Segal, 2002).

In this study, we investigated the effects of cholinergic antagonists with different selectivities to determine the receptors most responsible for the DBS-related increases in TrkA. We show that administration of mecamylamine but not scopolamine attenuated the increased TrkA expression, which suggests that the nicotinic, and not muscarinic, receptor subtypes are necessary for these stimulation effects. Further experimentation with selective nicotinic antagonism with DHBE or MLA on α4β2 and α7 receptors respectively indicated that the augmentation of TrkA is dependent on activity of α7 receptors and not α4β2 receptors.

To further expand on this receptor’s importance, this study utilized a *CHRNA7*^*lox/lox*^ mouse model and conditional knock-out approach with viral vector delivery of Cre-GFP using a SYN promoter. This system selectively knocked-down α7 receptors specifically in neurons while avoiding effects on the glial population of α7 dependent pathways which have been shown to promote anti-inflammatory and antioxidant cascades (Parada et al., 2013). We assumed that fluorescent cortical tissue that expressed GFP achieved an appropriate level of knocked-out α7 receptors, though measurement of the degree of knock-down was not performed. The saline control in the contralateral hemisphere also has limitations. A SYN-promoting viral vector that delivers GFP without Cre would better control for any possible effects of targeted immune response. Our results showed a decline in TrkA in the fluorescent tissue compared to the contralateral hemisphere, and this decline occurred in stimulated or unstimulated mice. These results suggest that α7 receptor activation is a necessary step for the maintenance of TrkA expression, and hence cholinergic axons, in the neocortex, which has previously been shown to be reliant upon the constitutive expression of NGF (Debeir et al., 1999). The relative equivalence of the effect in stimulated and unstimulated animals suggests that these pathways are under some level of activation on a near daily basis.

The potential mechanism of the α7 receptor’s role in cholinergic maintenance may stem from its facilitation of both ionotropic and metabotropic signaling. During its activation, structural changes result in conductance of Ca2+ through a transmembrane pore, which contributes to short term depolarization events. Its addition, G-protein Gαq subunit also interacts with phospholipase C and inositol-triphosphate to release Ca2+ from the endoplasmic reticulum (Kabbani & Nichols, 2018). This dual signaling with divalent cations results in significantly larger and longer periods of depolarization. This stronger divalent signaling may influence the release of proNGF and tissue plasminogen activator, tPA.

Previous studies demonstrate that postsynaptic vesicles of proNGF are released into the synapse constitutively and during low frequency firing of neuronal activity. Their interaction with p75NTR promotes cell death and axonal withdrawal (Song et al., 2010; Woo et al., 2005). During periods of high frequency activity, serine protease cleavage of neurotrophins, presumably by tPA, also occurs (Nagappan et al., 2009). Furthermore, the prolonged activation of the neocortex by cholinergic agonists results in the release of proNGF and tPA from the same neurons (Bruno & Cuello, 2006). The predominant neurons that release proNGF are parvalbumin containing interneurons (Biane et al., 2014). It stands to reason that the increased availability of tPA in synapses receiving high frequency action potentials would also lead to increased cleavage of proNGF to mNGF, as it is released in synapses constitutively and shares the same cleavage enzyme as proBDNF. This would explain the increases in both TrkA and TrkB signaling from forebrain stimulation that we have been previously demonstrated (Kumro 2023). Some neocortical interneurons activated by acetylcholine are sensitive to α7 nicotinic antagonists, suggesting they express α7 receptors(Alkondon et al., 2000). The α7 nicotinic receptor’s repeated activation by intermittent DBS hypothetically would cause more high frequency periods of neuronal activity, which would release both proNGF and tPA, which would enable tPA-dependent pathways to cleave proNGF into mNGF and activate the TrkA receptor.

The necessity of the α7 receptor in cholinergic neuron maintenance fits with previous literature. Long-term α7 receptor activation increases receptor density and neuroprotective mechanisms as well as improves cognition in multiple animal models (Bitner et al., 2010; Jonnala et al., 2002; Jonnala & Buccafusco, 2001). Allosteric modulators of α7 receptors have been shown to augment learning and memory in rodents and nonhuman primates when combined with cholinesterase inhibitors (Callahan et al., 2013). With the α7 receptor’s high binding affinity to and antagonism by amyloid-beta (H. Y. Wang et al., 2000), the cognitive decline associated with increased amyloid-beta in AD may be caused by beta-amyloid oligomers impairing necessary signaling for adequate maintenance of the cholinergic system similar to the pharmacological effects in this study. This notion is further supported by studies which showed loss of α7 receptors enhances amyloid-beta accumulation and cognitive decline in mice (Hernandez et al., 2010) as well as its reported protective role in AD (M. Hernandez & T. Dineley, 2012).

As prior work demonstrated, basal forebrain stimulation also reduced Aβ_42_ and BACE1, the rate limiting step in formation of Aβ_42_, the same step-wise pharmacological approach was used to determine acetylcholine receptor necessity for BACE1 reduction. Unlike the pronounced blunting effects of various antagonists when measuring TrkA expression, only a mild blunting of the stimulation-related reduction in BACE expression was observed after non-selective antagonist administration. This difference suggests a more indirect or redundant mechanism involving acetylcholine transmission and BACE1 expression. BACE1 has notable influences by multiple receptors. Its interaction with M1 muscarinic receptors mediates its proteosomal degradation (Jiang et al., 2012; Züchner et al., 2004). Additionally, p75NTR activation signals ceramide activation and BACE1 stability (Dobrowsky & Carter, 1998; Han et al., 2002; Venable et al., 1995). Normally the increased acetylcholine release from basal forebrain stimulation should contribute to both of these effects directly by stimulating M1 receptors and also promoting cleavage of proNGF to mNGF resulting in less p75NTR activation. The administration of either mecamylamine or scopolamine would only antagonize one of these pathways leaving the other unchecked, so it is plausible that BACE1 levels would still decrease albeit in a less pronounced fashion which is what we observed. A combined mecamlyamine/scopolamine administration should result in stimulation no longer suppressing BACE1 expression.

In summary, our study shows that basal forebrain DBS both augments TrkA expression and reduces BACE1 levels. We show that the TrkA pathway is uniquely dependent upon activation of α7-nACh receptors, while BACE1 expression is more loosely associated with specific cholinergic receptors. This research supports previous studies that show the importance of α7-nACh receptors in cognition and early AD pathology and offers a potential treatment for preventing the loss of cholinergic neurons as well as decreases the BACE1 dependent production of beta-amyloid which are both major processes that contribute to the onset and neuropathology of AD.

## Acknowledgments

Founders for the 5xFAD mice were graciously provided by Raghavan Raju. Kendyl Pennington assisted JK in stimulation of the mice.

## Author contributions

DTB and JK and AP designed all studies. JK performed all surgeries and all brain stimulation for behavioral and pharmacological experiments. ATerry designed all pharmacological dosing. JK and ATripathi performed all biochemical analyses in AP lab. DTB and JK drafted the manuscript with feedback from all authors.

Correspondence to David Blake

## Supplement

Supplementary data consists of original blots from Figures 1 through 4. The letter S will be added to the Figure name, and it will otherwise correspond to the primary figure in the manuscript e.g., Figure S1 will contain original blots from Figure 1. S5 and S6 contain the additional samples 4 and 5 of each group and were used in figures corresponding to their specific pharmacologic group. Dotted-line boxes indicate representative lanes used for their respective figure.

**Figure S1.**
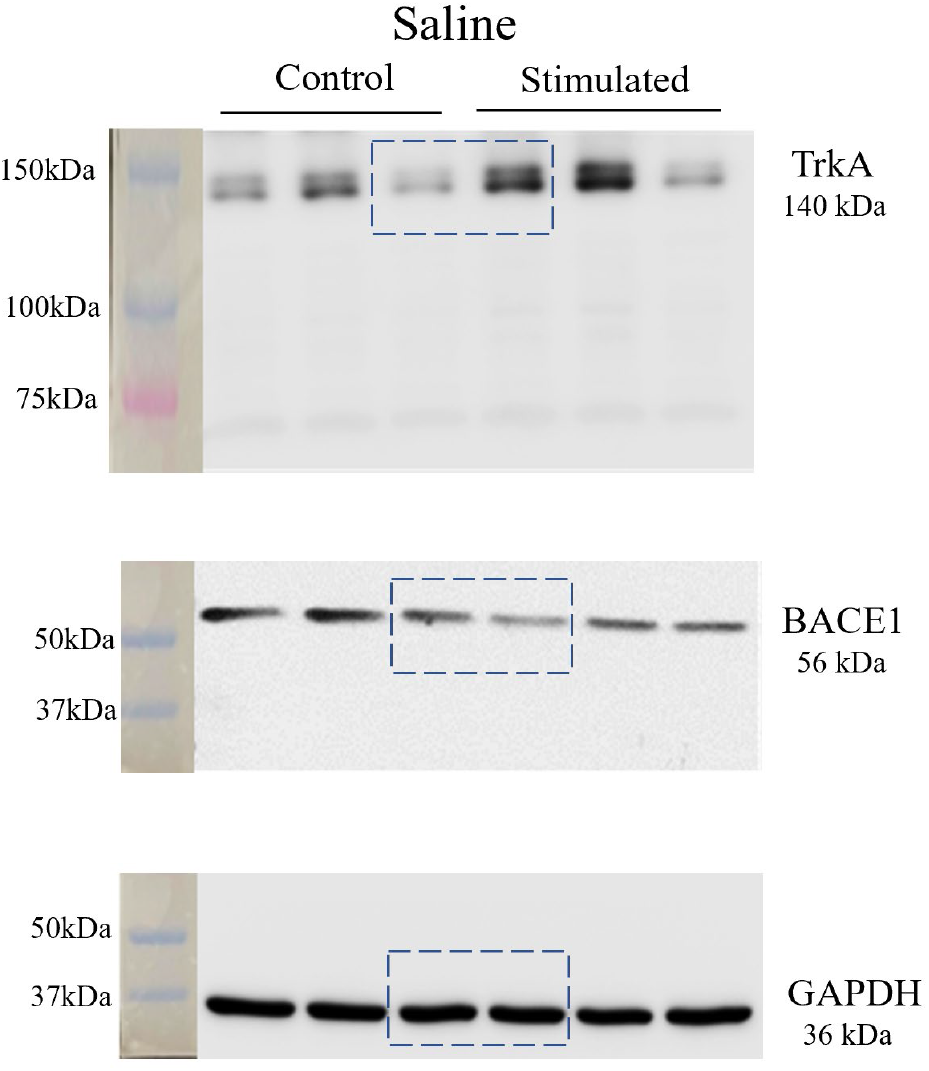

**Figure S2.**
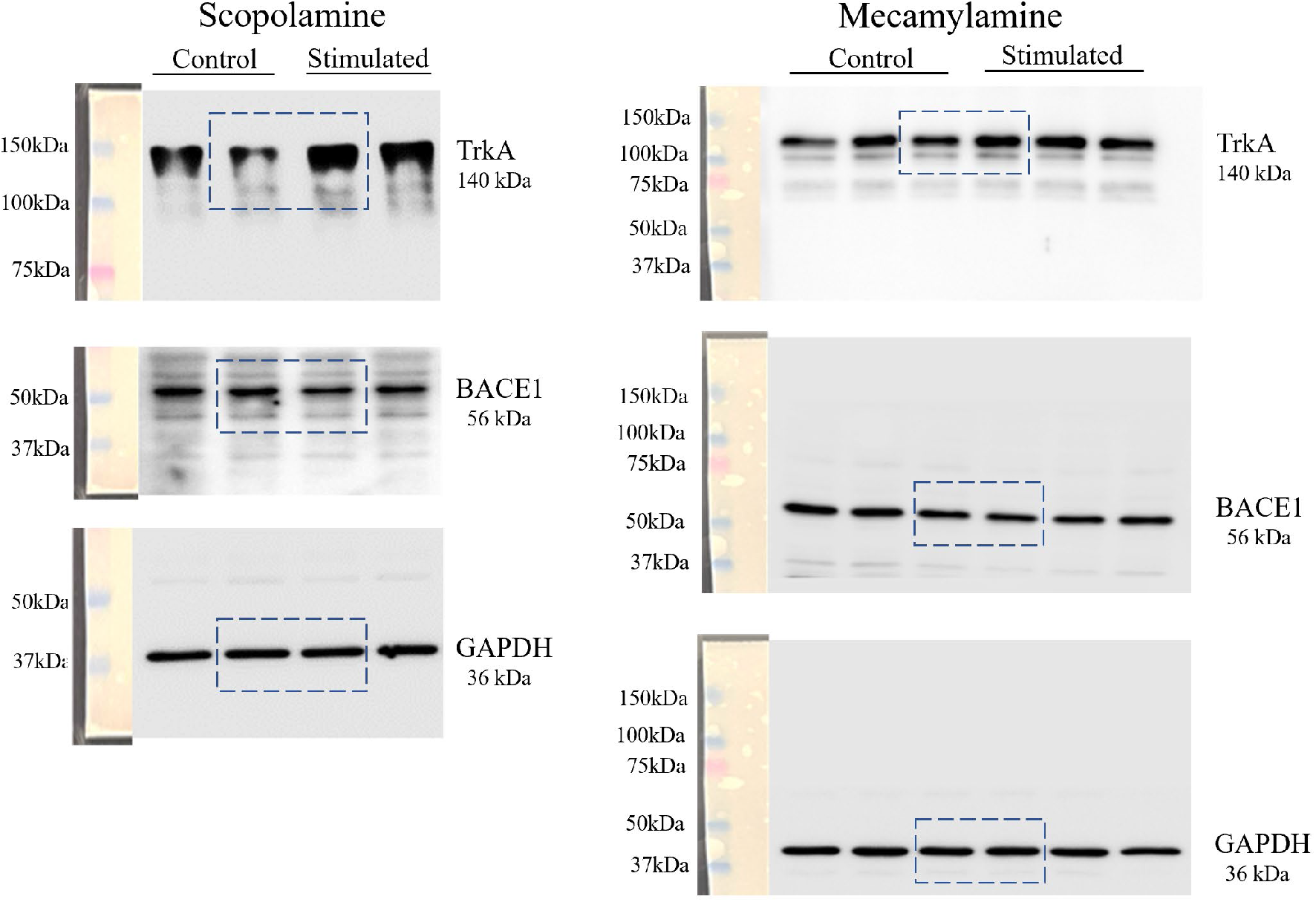

**Figure S3.**
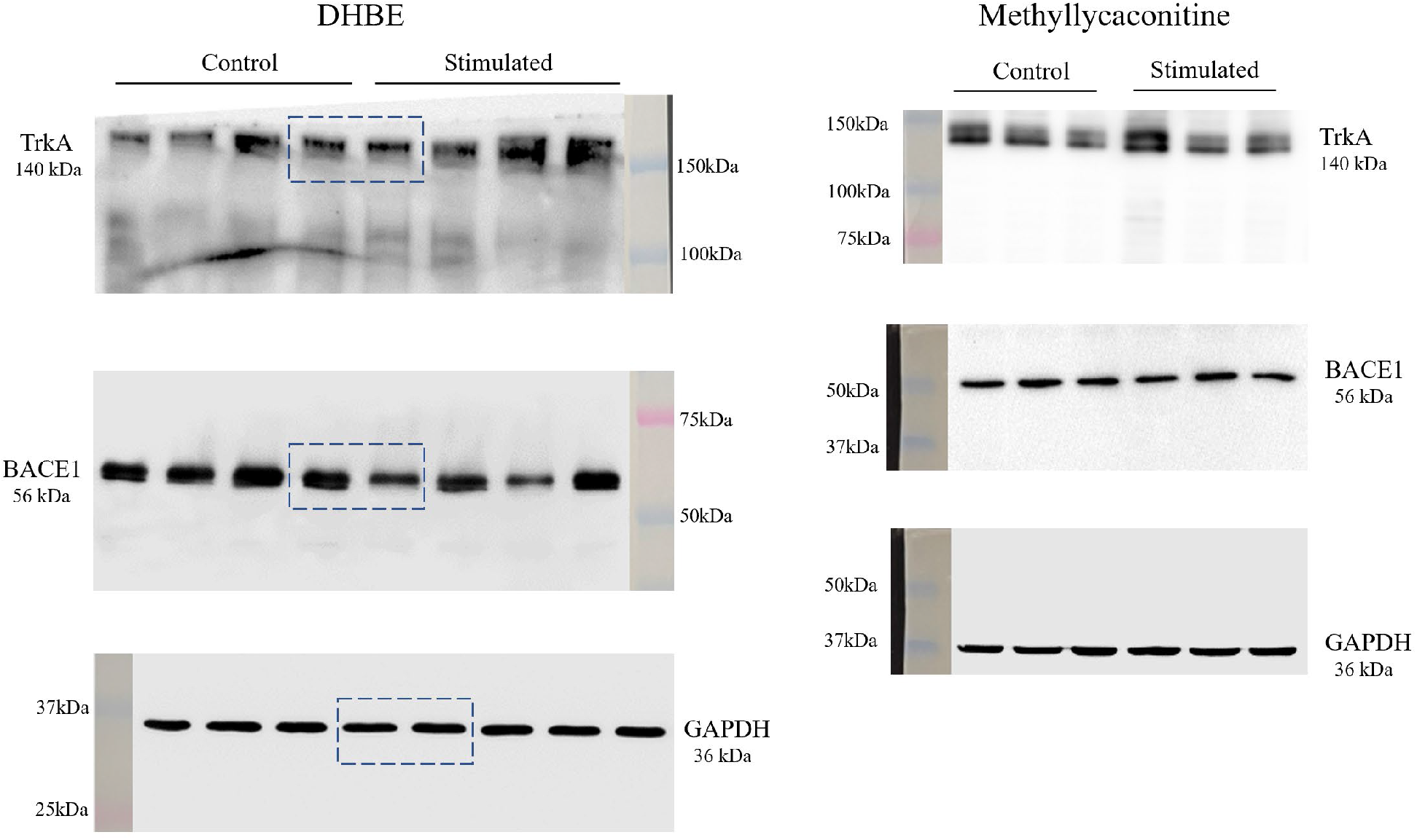

**Figure S4.**
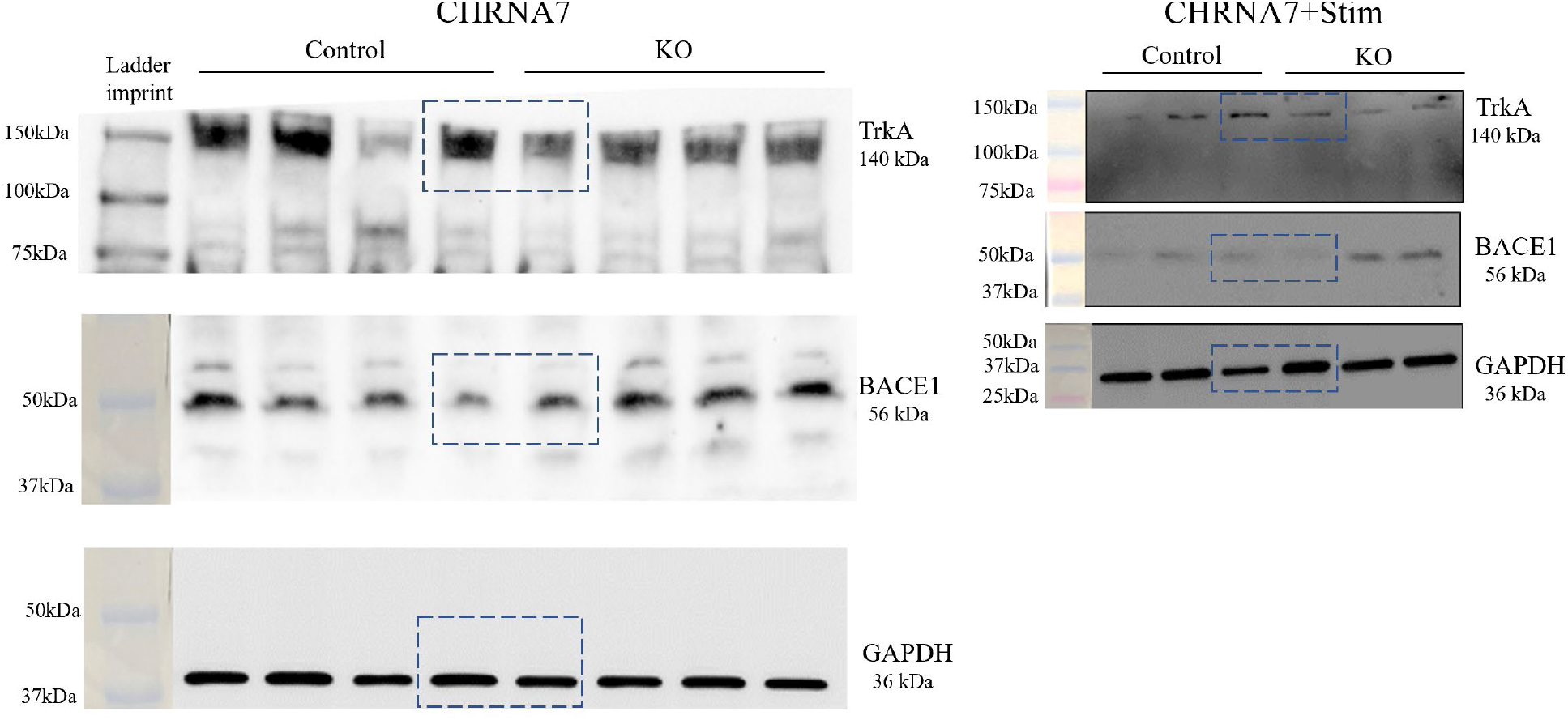

**Figure S5.**
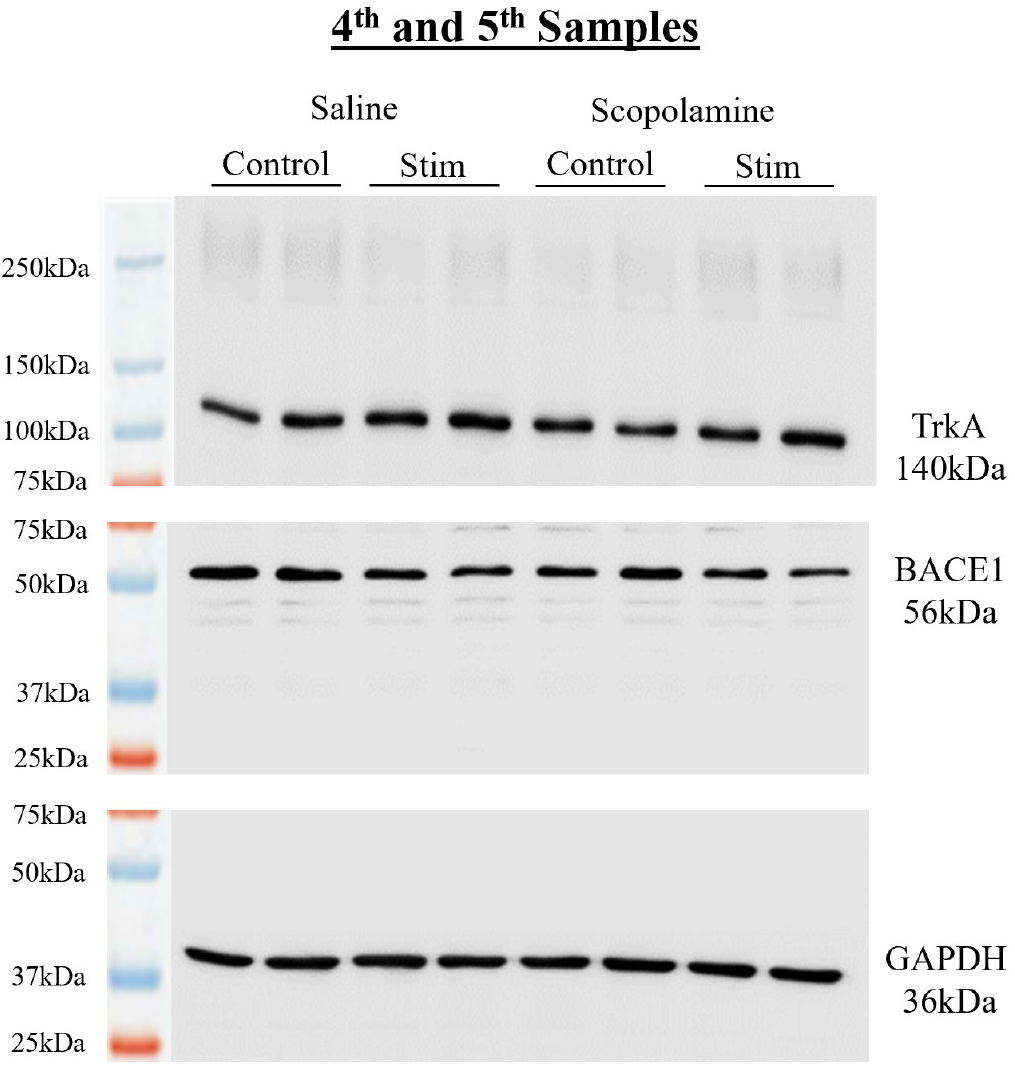

**Figure S6.**
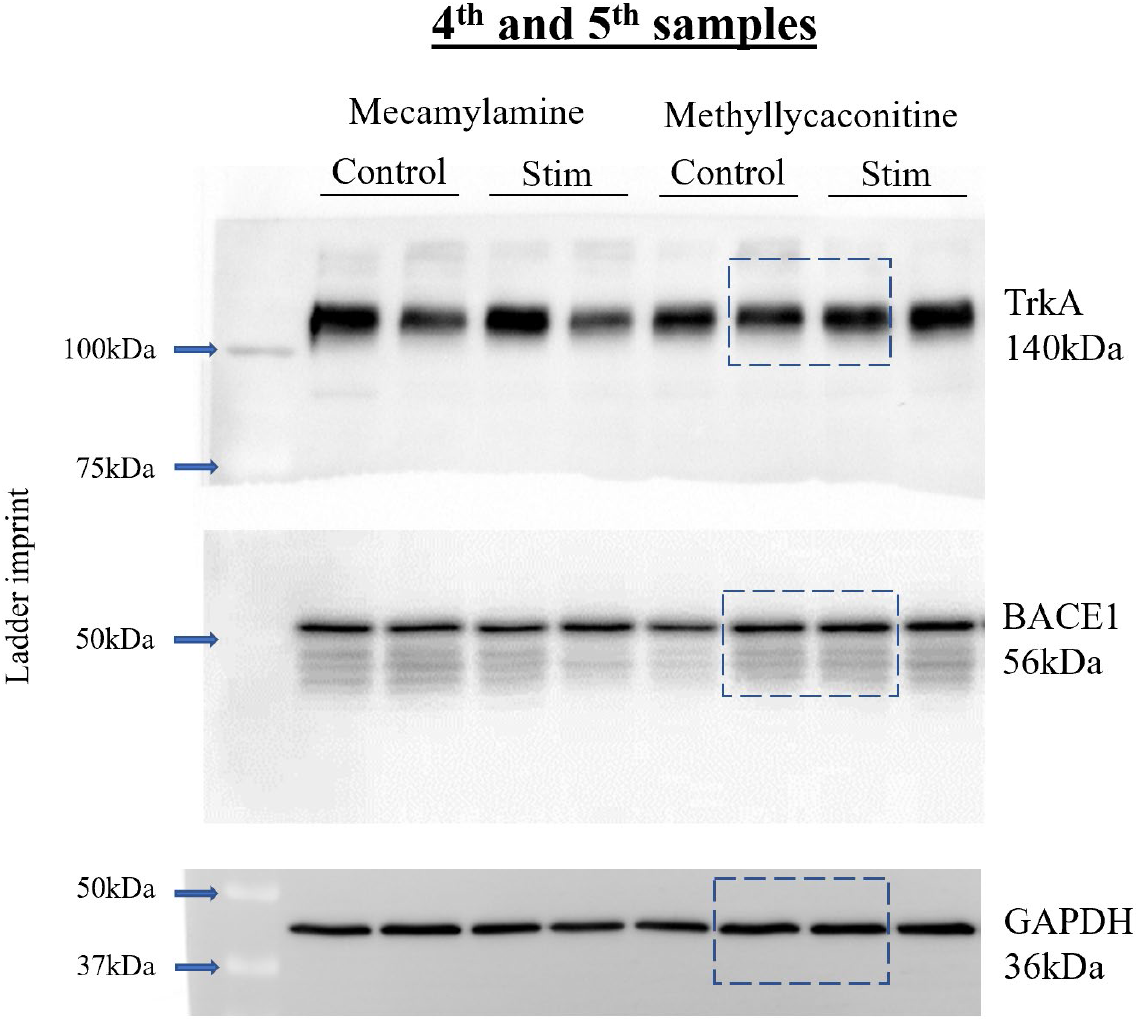

